# Genetic studies with a uracil DNA glycosylase biosensor support a role for mitochondrial UNG1 in nuclear uracil repair

**DOI:** 10.64898/2026.04.30.721890

**Authors:** Yu-Hsiu T. Lin, Analisa M. Lott, Xingyu Liu, Layla Abdulbaki, Yanjun Chen, Michael A. Carpenter, Reuben S. Harris

**Affiliations:** Department of Biochemistry and Structural Biology, University of Texas Health San Antonio, San Antonio, TX 78229, USA; Howard Hughes Medical Institute, University of Texas Health San Antonio, San Antonio, TX 78229, USA

**Keywords:** C-to-U deamination, cytosine deamination, uracil base excision repair (UBER), uracil-DNA N-glycosylase (UNG), uracil glycosylase inhibitor (Ugi)

## Abstract

The universally conserved enzyme uracil-DNA *N*-glycosylase (UNG) plays a central role in maintaining genome stability. It functions as the initiating factor in uracil base excision repair (UBER) by catalyzing the removal of uracil lesions in genomic DNA, a necessary first step in restoring genome integrity after hydrolytic deamination of cytosine to uracil or misincorporation of deoxy-uridine monophosphate during replication. Although methods have been developed to study UBER *in vitro* and *in cellulo*, none provide a quantitative readout of UNG activity on the chromosomal DNA of living cells. To address this gap, we created an UNG biosensor (U-report) that utilizes a modified cytosine base editor to generate a targeted genomic uracil lesion in a fluorescent reporter for C-to-U editing activity. UNG activity ablation through uracil DNA glycosylase inhibition (Ugi) or *UNG*-knockout results in elevated reporter fluorescence. Isoform-specific knockouts show that mitochondrial UNG1 also contributes to UBER of nuclear DNA. Our studies establish a real-time biosensor for quantification of chromosomal DNA uracil excision activity in living cells and, in support of prior studies, indicate that both UNG isoforms should be addressed in small molecule inhibitor development programs.

**Graphical abstract:** 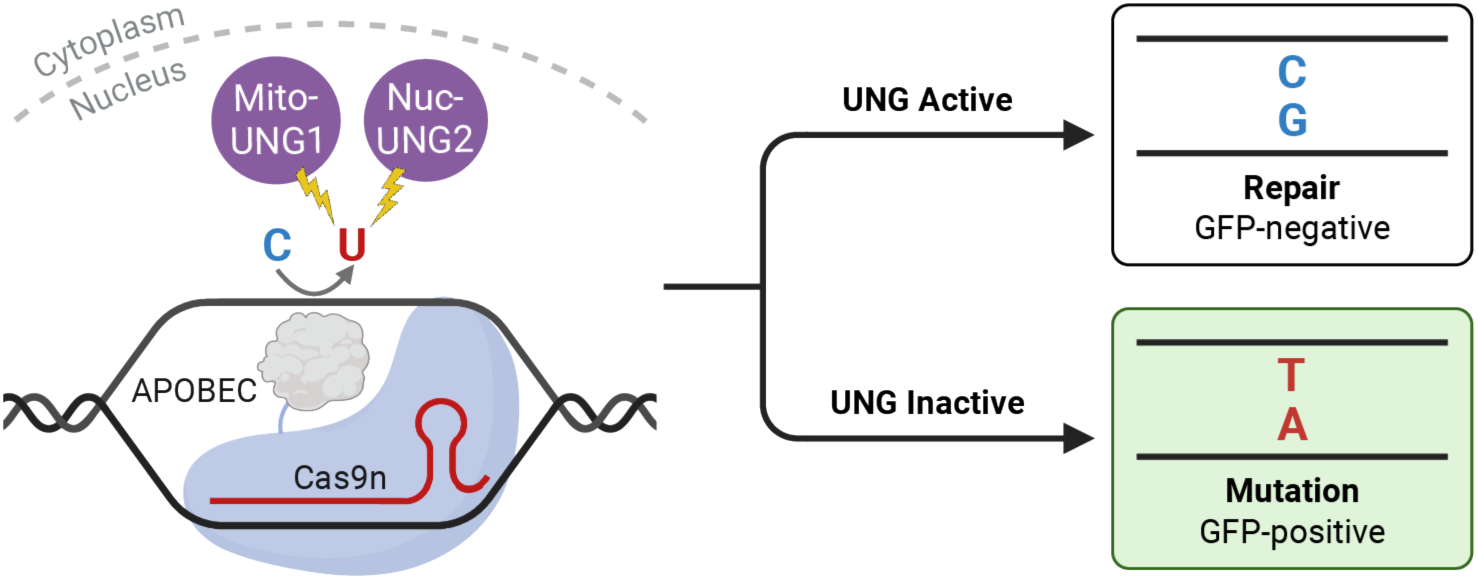

## Introduction

Uracil (U) is often regarded as the fifth base, after the canonical nucleobases adenine (A), cytosine (C), guanine (G), and thymine (T). Like T, U forms two hydrogen bonds with A and, following the Central Dogma, is one of the defining features of RNA polynucleotides templated from DNA substrates. However, uracil is chemically distinct from thymine, as it has a hydrogen instead of a methyl group at the 5-position of the pyrimidine ring structure. Uracil exists in several forms in cells including as a free base, as a nucleoside coupled to a ribose sugar, and as mono-, di-, and tri-phosphorylated nucleotides (UTP being the most abundant) [1,2]. Uracil can also be altered chemically by several different RNA modification enzymes, predominantly in the context of RNA polynucleotides (*e.g*., site-specific pseudo-uridine modifications in rRNAs and tRNAs) [3]. These modifications are less abundant in cells but add important layers of epigenetic information and structural features [4].

Although integral in RNA polynucleotides, uracil is a non-canonical DNA base. It can be misincorporated into DNA during replication by escaping the DNA polymerase base selection and proofreading mechanisms [5]. Most of these replication-associated uracils pair faithfully with adenines and are removed and replaced faithfully with thymine by subsequent uracil base excision repair (UBER, described in greater detail below). Uracils can also occur in DNA following spontaneous and enzyme-catalyzed hydrolytic deamination of cytosine to uracil. Activation-induced cytidine deaminase (AID), apolipoprotein B mRNA editing enzyme, catalytic polypeptide-like 1 (APOBEC1), and seven different APOBEC3 enzymes (A3A-D, F-H) in human cells catalyze the deamination of C-to-U in single-stranded (ss)DNA substrates. This enzymatic reaction is physiologically important for antibody gene diversification by somatic hypermutation and class switch recombination (AID) and for innate antiviral immunity (A3s and possibly also AID and APOBEC1) [6–8]. These DNA uracils can also be corrected by UBER, but in specific contexts, such as R-loops or recombination intermediates, the U:G mismatches resulting from deamination may be recognized and processed by other DNA repair pathways including mismatch repair [9–11].

UBER is a universally conserved and mostly error-free mechanism that serves to remove uracil from double-stranded DNA substrates [5,12]. The first step in this DNA repair pathway is uracil recognition and excision by uracil-DNA *N*-glycosylase (UNG). Mammalian cells express two isoforms through alternative transcription initiation and alternative splicing upstream of shared exons encoding a conserved UNG domain [13]. UNG1 has a unique mitochondrial localization sequence (MLS) for UBER in mitochondria, whereas UNG2 has a nuclear localization signal (NLS) that promotes localization to the nucleus and excision of chromosomal DNA uracils. Following the generation of an apyrimidinic site, the second step is cleavage of the sugar-phosphate backbone by the apurinic-apyrimidinic endonuclease APE1. This produces a single-stranded break with a 3’-OH and a 5’-deoxyribose phosphate (dRP). The third step is removal of the 5’-dRP and templated incorporation of a single nucleobase, usually error-free, by DNA polymerase β (POLβ). The fourth and final step is ligation of the phosphodiester backbone by a DNA ligase (usually LIGIII) to restore the integrity of the DNA helix.

Interestingly, UBER is potently inhibited by proteins that bind to UNG and block its activity by mimicking DNA substrates [14]. The *Bacillus subtilis* bacteriophages PBS1 and PBS2 have evolved to utilize uracil instead of thymine in their genomic DNA and, accordingly, these phages encode a small 9.5-kDa protein called uracil-DNA glycosylase inhibitor (Ugi) that mimics the structure of a flipped-out deoxy-uridine nucleotide [15]. This leads to a near-irreversible interaction between UNG and Ugi with potent inhibition of UNG at equimolar concentrations [15]. In addition, the *B. subtilis* phage φ29, a non-uracil-containing DNA virus, encodes p56 that dimerizes to inhibit bacterial UNG [16,17]. Furthermore, *Staphylococcus aureus* encodes a uracil-DNA glycosylase inhibitor that has a high affinity to its own *S. aureus* uracil-DNA glycosylase [18]. Because of the high degree of conservation in the mechanism of action of Ugi and Ugi-like proteins, as well as UNG, across the kingdoms of life, these protein inhibitors can function against a broad range of UNG enzymes from multiple species including humans (*e.g*., [19,20]).

The biological importance of UNG in UBER, as exemplified by its presence in multiple kingdoms of life and the evolution of counteraction mechanisms in bacteria and viruses, has motivated efforts to measure its activity in both biochemical and cellular contexts. The first enzymatic assay was developed by Tomas Lindahl to monitor the release of radioactively labeled uracils from polydeoxynucleotides by paper chromatography [21]. Other classical methods detect the cleavage of a radioactively or fluorescently labeled deoxyuridine-containing DNA (either single- or double-stranded) by gel electrophoresis [22–24]. Moreover, approaches using molecular beacons with fluorogenic properties have also been established to enable real-time reporting both *in vitro* and in cells [25–31]. Notably, a cell-based assay utilizes a BFP reporter whose expression is sensitive to the recognition and repair of a pre-installed U:G base pair [32]. Though many tools are available for assessing UNG activity, they are either incompatible with live cells or do not directly measure excision of uracils in chromosomal DNA, which are thought to be the predominant substrates of UNG2.

Here, we report a biosensor called U-report for detection and quantification of UNG-catalyzed excision of a chromosomal DNA uracil generated *de novo* in living cells. This assay utilizes a modified cytosine base editor to generate a targeted uracil lesion in an *eGFP* reporter. Faithful UBER prevents *eGFP* reversion and results in low fluorescence, whereas UNG inhibition through Ugi expression or CRISPR-mediated gene inactivation results in uracil persistence and high fluorescence. This U-report system is effective in both episomal and chromosomally integrated formats. Moreover, the systematic ablation of the nuclear UNG2 and mitochondrial UNG1 proteins showed that the latter enzyme is also capable of functioning in UBER in the nuclear compartment of human cells.

## Materials and methods

### Cell lines

Cells were maintained in a humidified incubator (Fisher Scientific 11-676-600) at 37°C and 5% CO_2_. 293T cells (ATCC CRL-3216) were cultured in DMEM, high glucose media (Thermo Fisher Scientific 11965092) supplemented with 10% fetal bovine serum (Biowest 058N24) and 1X penicillin-streptomycin (Thermo Fisher Scientific 15140122). U2OS cells (ATCC HTB-96) were grown in McCoy’s 5A media (Thermo Fisher Scientific 16600082) supplemented with 10% fetal bovine serum and 1X penicillin-streptomycin. Knockout cell lines were generated by delivering the ribonucleoprotein (RNP) complex (Cas9 nuclease from Thermo Fisher Scientific A36497 and custom sgRNA from Invitrogen TrueGuide Synthetic gRNA) using the Lipofectamine™ CRISPRMAX™ Cas9 Transfection Reagent (Thermo Fisher Scientific CMAX00001). Guide RNAs targeting *UNG1* (AGCTTCCGGCCCAACCCCCA), *UNG2* (CGTCTTCTGGCCGATCATCC), and *UNG1/2* (GCGGCCCGCAACGTGCCCGT) were used. Briefly, 90,000 293T and 50,000 U2OS cells were seeded in a 24-well plate (Corning 3527) and transfected the following day with the RNP complex according to the manufacturer’s protocol. After 24 hours, cells were treated with trypsin (Thermo Fisher Scientific 25300120), counted, and seeded at 0.5 cells per well in 96-well plates (Corning 3596). Isogenic clones were expanded and screened by immunoblotting and uracil excision assay (details below). Insertion-deletion mutations (indels) at the targeted loci were confirmed by long-read amplicon sequencing using Oxford Nanopore Technology (Standard Genotyping Analysis, Plasmidsaurus) and by Sanger sequencing of ten individual amplicons cloned into a pJET1.2 vector (Thermo Scientific K1232). Genotypes were manually annotated for coding changes (c.) and corresponding protein-level effects (p.) using the HGVS nomenclature. Specifically, the p.0 annotation indicates disruption of the ATG start codon and no protein production.

### Reporter constructs

The parental A3B^ctd^-BE4max plasmid with two tethered Ugi proteins (AddGene #198888), the Y93H TC AMBER reporter construct (AddGene #198882), and the Y93H TC AMBER gRNA (AddGene #256530) are available from AddGene (https://www.addgene.org/search/catalog/plasmids/?q=reuben+harris) [33]. Sequences encoding Ugi proteins were removed from A3B^ctd^-BE4max by site-directed mutagenesis (SDM) to generate A3B^ctd^-BE4max^ΔUgi^ (AddGene #254664, pRH11239). Ugi was subcloned from the A3B^ctd^-BE4max construct into a pcDNA3.1 backbone by In-Fusion Snap Assembly (Takara Bio 638947). A Ugi-FLAG construct (AddGene #254665, pRH11253) was subsequently generated by SDM. The AMBER fluorescent system was cloned into a pCMV backbone by In-Fusion for transient transfection experiments. The human UNG2 sequence (NM_080911.3) was synthesized as a gBlock (Integrated DNA Technologies) and cloned into a pcDNA3.1 backbone (AddGene #254663, pRH11236). **Supplementary Table S1** provides a list of all oligonucleotides used for cloning and sequencing.

### Transient transfection experiments

150,000 293T and 60,000 U2OS cells per well in 1 mL complete medium were seeded in 12-well plates (Corning 3512) and allowed to adhere overnight. Transfection was then performed with 100 ng of each construct (CBE, gRNA, reporter, Ugi) and supplemented with pcDNA3.1 empty vector as needed to maintain the same total DNA amounts for all reactions. A 1:3 ratio of plasmid (μg) to *Trans*IT-2020 (Mirus Bio MIR 5405) transfection reagent (μL) was incubated in 100 μL Opti-MEM (Thermo Fisher Scientific 31985088) for 15 mins at room temperature (RT) before transferring the mixture drop-wise to cells. A master mix was prepared for each condition and split across three wells for technical replicates. After transfection, cells were placed in a humidified incubator at 37 °C for real-time imaging by Sartorius Incucyte SX5 followed by endpoint analyses by flow cytometry and immunoblotting.

### Chromosomal experiments

Stable 293T cell lines expressing the AMBER system were generated by lentiviral transduction, single cell cloning, and screening for uniformly high mCherry signal. Lentiviral stocks were produced by transfecting 293T cells with the Y93H TC AMBER reporter plasmid (AddGene #198882, pRH9898) [33], a packaging plasmid, and a vector encoding the VSV-G envelope protein. Viral supernatants collected at 48 and 72 hours post-transfection were combined and passed through a 0.45 μm PVDF filter (Millipore SLHVR33RB). To generate a single- or low-copy 293T cells with a chromosomally integrated AMBER reporter, cells were infected at low multiplicity of infection (<0.01) and single-cell sorted for mCherry-high expressors into 96-well plates using the BD FACSAria Fusion flow cytometer. Candidate mCherry-high clones were expanded and subjected to flow cytometry. The single clone with the strongest and most uniform mCherry expression was selected for downstream experimentation (293T-chromosomal AMBER c15 or 293T-cAMBER c15). Alu-PCR attempts to identify the AMBER insertion site(s) were not successful, but a uniformly high mCherry signal by flow cytometry of 293T-cAMBER c15 strongly indicates a single insertion site (**Supplementary Fig. S1**).

### Immunoblotting

Cell pellets were lysed in ice-cold RIPA buffer (50 mM Tris, 150 mM NaCl, 0.5% sodium deoxycholate, 1% NP-40, 0.1% SDS, pH 8.0) supplemented with 1X protease inhibitor cocktail (Roche 11836170001). Lysates were vortexed vigorously and sonicated in a water bath sonicator at 4 °C for 10 mins. Insoluble material was pelleted at top speed for 10 mins at 4 °C. Clarified lysates were transferred to a new tube, and protein concentrations were determined using a BCA Protein Assay (Thermo Scientific 23227). 20 μg of lysate was mixed with reducing Laemmli SDS sample buffer (Thermo Scientific J61337.AC) and boiled at 95 °C for 5 mins. Samples and protein ladder (Thermo Scientific 26619) were loaded onto a 4-20% TGX Stain-Free Precast Gel (Bio-Rad 5678094) and run in Tris-Glycine-SDS buffer (25 mM Tris base, 200 mM glycine, 0.1% SDS) at 90 V for 2 hours. Proteins were transferred to a PVDF FL membrane (Millipore Sigma IPFL00005) at 90 V for 2 hours on ice. Membranes were blocked in 1X casein buffer (Sigma-Aldrich B6429) for 1 hour at RT. Primary and secondary antibodies were incubated in blocking buffer at 4 °C overnight and at RT for 1 hour, respectively. Membranes were washed four times at RT for 5 mins each on shaker after each antibody incubation. Primary antibodies include 1:1,000 α-Cas9 (CST 14697S, lot 8), 50 ng/mL α-UNG (CST F8Z1L, lot VSP160679-1), 1:1,000 FLAG (Sigma-Aldrich F1804, lot 315235), 1:5,000 α-alpha-tubulin (Sigma-Aldrich T5168, lot 165669). Secondary antibodies include 1:5,000 IRDye 800CW Goat α-Mouse (LICOR 926-32210, lot D40618-15) and 1:10,000 IRDye 680RD Goat α-Rabbit (LICOR 926-68071, lot D40109-05). Immunoblots were imaged using a LI-COR Odyssey M imaging system.

### Fluorescence microscopy

Live-cell images were captured using a Sartorius Incucyte SX5 instrument. Nine images were acquired per well in one-hour increments from the time of transfection to sample collection for flow analysis. Data were collected in the phase, green (300 millisecond acquisition time), and orange (400 ms acquisition time) channels using the 20X objective and standard scan type. Total integrated intensity for eGFP and mCherry was quantified in the green and orange channels, respectively, using the Incucyte Basic Analysis Software. To control for cell confluency, normalized integrated intensity was calculated by dividing the total integrated intensity by the plate area occupied by cells in the phase channel. For each well, the mean quantification was taken across the nine images.

### Flow cytometry

Cells were harvested by washing with PBS followed by trypsinization. Single-cell suspensions were loaded into a 96-well U-bottom plate (Corning 353077) for data acquisition on a BD LSRFortessa X-20 flow cytometer coupled to a BD High Throughput Sampler. For every sample, 30,000 singlet events were analyzed. Downstream flow analysis and calculations of median fluorescence intensity were conducted using FlowJo v10.10.1.

### Uracil excision assay

The *in vitro* uracil excision assay was performed as described [11]. Briefly, cell pellets were lysed with HED buffer (25 mM HEPES, 15 mM EDTA, pH 7.4) by four freeze-thaw cycles in liquid nitrogen and 37 °C water bath. Insoluble material was pelleted at top speed for 10 mins at 4 °C. Clarified lysates were transferred to a new tube, and protein concentrations were determined using the Bradford reagent (Sigma-Aldrich B6916). Reaction volume of 50 μL containing 20 pmol of fluorescent oligo 5’-ATTATTATTATT<u>U</u>TAATGGATTTATTTATTTATTTATTTATTT-6-FAM and normalized amounts of lysate was prepared in HED buffer. Negative and positive controls with HED buffer and recombinant *E. coli* uracil-DNA glycosylase (New England Biolabs M0280S, lot 10272153), respectively, were included. Reactions were incubated at 37 °C for 10 mins followed by inactivation at 95 °C for 5 mins. Cleavage at abasic sites was induced by adding 5 μL 1M NaOH and incubating at 95 °C for 5 mins. Prior to loading 1 pmol of the substrate to a 20% TBE-Urea gel, samples were mixed with an equal volume of loading buffer containing 80% formamide and heated at 95 °C for 5 mins. Gels were run in 1X TBE buffer at 10 watts for 35 mins and imaged on a LI-COR Odyssey M system.

### Statistical analyses

Data analyses and visualization were performed in R version 4.5.2. Welch’s two-sample two-sided *t*-tests were conducted to determine statistical differences between conditions. Unless otherwise stated, statistical significance is indicated as n.s. = not significant; \**P*< 0.05; \*\**P* < 0.01; \*\*\**P* < 0.001.

## Results

### UNG biosensor design and validation (U-report system)

To monitor UNG activity in live cells, we built upon two prior advances – cytosine base editing (CBE) and APOBEC-mediated base editing reporter (AMBER) technologies. CBE technology utilizes a Cas9 nickase (Cas9n) complex to target a tethered APOBEC enzyme to a specific DNA cytosine [20,34] (**Fig. 1A**). This promotes efficient conversion of the target cytosine to uracil, which is protected from nuclear UNG excision by Ugi proteins tethered *in cis* to the same complex (usually two Ugi proteins). The resulting uracil is subsequently converted into a mutation by local DNA repair synthesis on the nicked and unedited DNA strand. We hypothesized that an UNG biosensor could be created by removing the Ugi proteins from CBE complexes (**Fig. 1B**). This strategy is predicted to enable cellular UNG to efficiently excise DNA uracils and trigger error-free UBER.

**Fig. 1.**
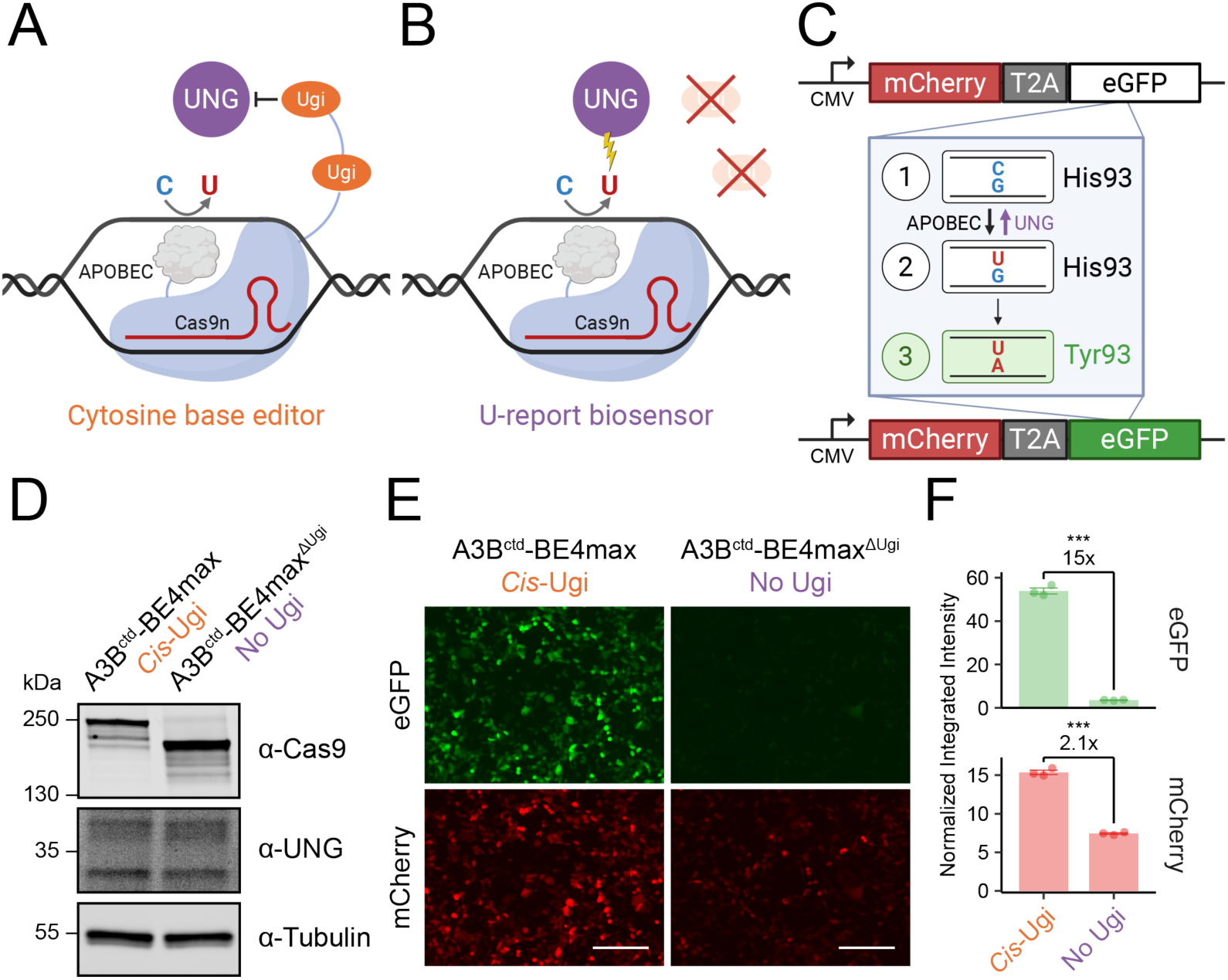
Engineering a biosensor to quantify UNG activity in living cells. **A)** Schematic of a current cytosine base editing technology. **B)** Schematic of the cellular UNG biosensor (U-report system) without UNG-inhibitory Ugi proteins. **C)** Schematic of the AMBER reporter for quantifying specific C-to-T mutations in eGFP. Steps 1 and 2 illustrate APOBEC-catalyzed deamination of the target cytosine. Steps 2 and 3 show the reactivation of eGFP activity by immortalizing the uracil lesion into a mutation and converting the transcribed (bottom) strand of dead eGFP back to a wildtype sequence. Importantly, UNG reverts step 2 back to step 1 (purple arrow) by excising the uracil and initiating error-free repair. **D)** Immunoblot showing expression of Cas9 and UNG in 293T cells 48 hours post-transfection. **E-F**) Live-cell fluorescence imaging and quantification of eGFP and mCherry signal under the indicated conditions at 48 hours post-transfection (scale=200μm; *n*=3, mean±SEM).

To monitor the progression of base editing, AMBER reporter technology was used for real-time quantification of CBE activity in living cells [33,35,36]. Precise DNA C-to-U editing of the AMBER construct causes reversion of a null allele of *eGFP* and the restoration of green fluorescence (**Fig. 1C**). Briefly, AMBER is a bicistronic DNA construct encoding mCherry, which is constitutively expressed, and eGFP, which is conditionally expressed. The fluorescent mCherry protein serves as an internal expression control. In the AMBER system, the eGFP reporter initially encodes a histidine at residue 93 that renders the protein non-fluorescent. Upon C-to-U editing and creation of a C-to-T mutation, His93 (<u>C</u>AC) is replaced by Tyr93 (<u>T</u>AC), which restores eGFP function. In detail, deamination of the target cytosine first produces a U:G mismatch (step 1 to 2 in **Fig. 1C**). How this mispairing is resolved ultimately dictates eGFP activity. Replication over the uracil or processing by mismatch repair (MMR) factors using the edited strand as the template will then convert the U:G into a U:A base pair, thereby restoring eGFP fluorescence (step 2 to 3 in **Fig. 1C**). Alternatively, if UNG removes the uracil, then base excision repair will revert the U:G back to C:G, leaving eGFP in an inactive state (step 2 to 1 in **Fig. 1C**). The net result is that UNG function will keep eGFP off, whereas UNG inhibition will help restore eGFP activity (illustrated in **Graphical abstract**). Taken together, by coupling the modified CBE lacking Ugi proteins (**Fig. 1B**) with AMBER (**Fig. 1C**), we envisioned that this strategy would generate a biosensor for detecting UNG activity and inhibition in living cells.

As a proof of concept, we first sought to test whether the assay is responsive to the activity of endogenous UNG in 293T cells. We compared a traditional CBE (A3B^ctd^-BE4max [33]), which has two Ugi proteins tethered *in cis*, to a modified version lacking Ugi (A3B^ctd^-BE4max^ΔUgi^; **Materials and methods**). These constructs were co-transfected into 293T cells along with plasmids for the Y93H TC AMBER gRNA and reporter, and the cells were analyzed 48 hours later by immunoblotting, fluorescence microscopy, and flow cytometry. The A3B^ctd^-BE4max^ΔUgi^ construct was expressed slightly better than the full CBE despite equal amounts of transfected plasmid, whereas the levels of endogenous UNG and α-tubulin were nearly identical (**Fig. 1D**). As expected, A3B^ctd^-BE4max with two tethered Ugi proteins promoted rapid and efficient AMBER editing and restoration of eGFP fluorescence (**Fig. 1E-F**). In contrast, despite higher steady state expression levels, the A3B^ctd^-BE4max^ΔUgi^ construct lacking Ugi promoted much lower levels of eGFP fluorescence, likely due to endogenous UNG counteracting base editing (**Fig. 1E-F**). A modest decrease in mCherry fluorescence was also observed in the absence of Ugi (**Fig. 1F**). Quantification of the normalized integrated intensity from live-cell fluorescence imaging indicates a 15- and 2.1-fold change in eGFP and mCherry levels, respectively (**Fig. 1F**). These changes in fluorescence were further supported by flow cytometry data (**Supplementary Fig. S2A-B**). Hereafter, we refer to the A3B^ctd^-BE4max^ΔUgi^ construct and the Y93H TC AMBER reporter as the U-report biosensor or the U-report system.

### U-report biosensor is sensitive to Ugi over-expression *in trans*

Using the U-report system, we next asked whether endogenous UNG can be inhibited by over-expression of Ugi *in trans*. We first confirmed equal expression of the modified CBE and UNG between the two conditions by immunoblotting (**Fig. 2A**). Upon introduction of Ugi by co-transfection, fluorescence microscopy showed a 15-fold increase in eGFP signal with an associated 1.9-fold increase in mCherry signal (**Fig. 2B**-**C**). Flow cytometry validates these findings by the pronounced shift in the double-positive population in the upward (elevated eGFP) and rightward (elevated mCherry) directions (**Fig. 2D**). Quantification of the median fluorescence intensity (MFI) from flow analysis indicates 14- and 1.7-fold changes in eGFP and mCherry activity, respectively (**Fig. 2E**). To assess the sensitivity of the U-report system, we co-transfected 0 to 300 ng of constructs encoding Ugi and human UNG2 in combination with the reporter plasmids and found that the biosensor is dose-responsive to a broad range of UNG activity levels (**Supplementary Fig. S3A-C**). Collectively, these results show that the U-report biosensor can measure the activities of both endogenous and exogenous UNG in 293T cells.

**Fig. 2.**
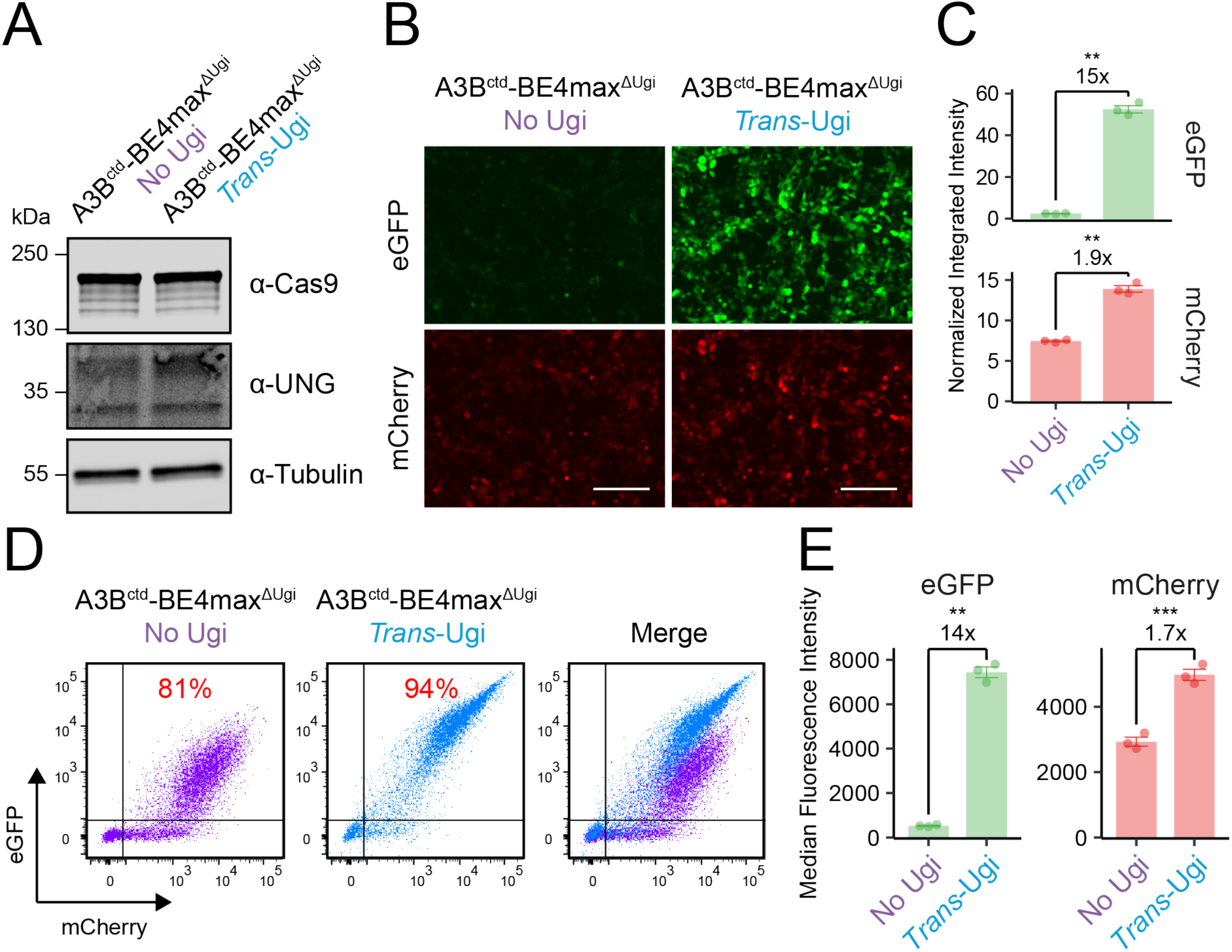
Ugi expression *in trans* inhibits UNG and produces strong eGFP signal. **A**) Immunoblot showing Cas9 and UNG levels in the tested conditions 48 hours post-transfection of 293T cells. **B-C**) Live-cell fluorescence imaging and quantification showing significant increases in eGFP and mCherry activity upon Ugi expression *in trans* at 48 hours post-transfection (scale=200μm; *n*=3, mean±SEM). **D-E**) Flow analysis and quantification of the singlet population at 48 hours post-transfection. The percentages in red indicate the ratio double-positive to mCherry-positive cell populations (*n*=3, mean±SEM).

### Chromosomal U-report biosensor stabilizes mCherry fluorescence levels

Although the increase in eGFP signal upon UNG inhibition is expected with the U-report system, the concomitant change in expression of the constitutive mCherry marker was unexpected, because it should have yielded similarly bright fluorescence readouts across the various conditions. A probable explanation for this observation is APOBEC-catalyzed restriction of foreign episomal DNA but not foreign integrated DNA [37–39]. This led us to hypothesize that stable expression of the AMBER reporter construct from chromosomal DNA may overcome the issue of variable mCherry expression, as well as generate a cleaner binary switch (*i.e*., an *eGFP* off-to-on switch). To test this idea, the AMBER reporter was delivered into 293T cells by lentiviral transduction with a low multiplicity of infection to ensure single- or low-copy integration events. Next, multiple clones were screened to identify a single clone (293T-chromosomal AMBER c15 or 293T-cAMBER c15) with strong and uniform mCherry expression (**Supplementary Fig. S1**). Last, constructs encoding CBEs, gRNA, and Ugi were co-transfected into 293T-cAMBER c15 cells for analysis 48 hours later. Expression of the CBEs and UNG were equal across all conditions (**Fig. 3A**). By fluorescence microscopy, we found that eGFP expression correlated positively with UNG activity through Ugi expression in *cis* or in *trans* (**Fig. 3B**) with fold changes of 44 and 51, respectively, relative to no Ugi (**Supplementary Fig. S4**). In addition, the mCherry signal was comparable across all conditions (<1.2-fold change) indicating greater stability with a chromosomal AMBER reporter system (**Supplementary Fig. S4**). These microscopy results are consistent with quantification by flow cytometry, which showed maximal fold changes in eGFP and mCherry MFI of 10.5 and 1.1, respectively, in the presence versus absence of Ugi (**Fig. 3C-D**). Together, these data indicated that stable integration of the AMBER reporter prevents variation in mCherry expression in response to changes in UNG activity levels. Importantly, these results also showed that the integrated U-report biosensor can quantify endogenous UNG activity on *de novo-*generated uracils in the context of chromosomal DNA of living cells.

**Fig. 3.**
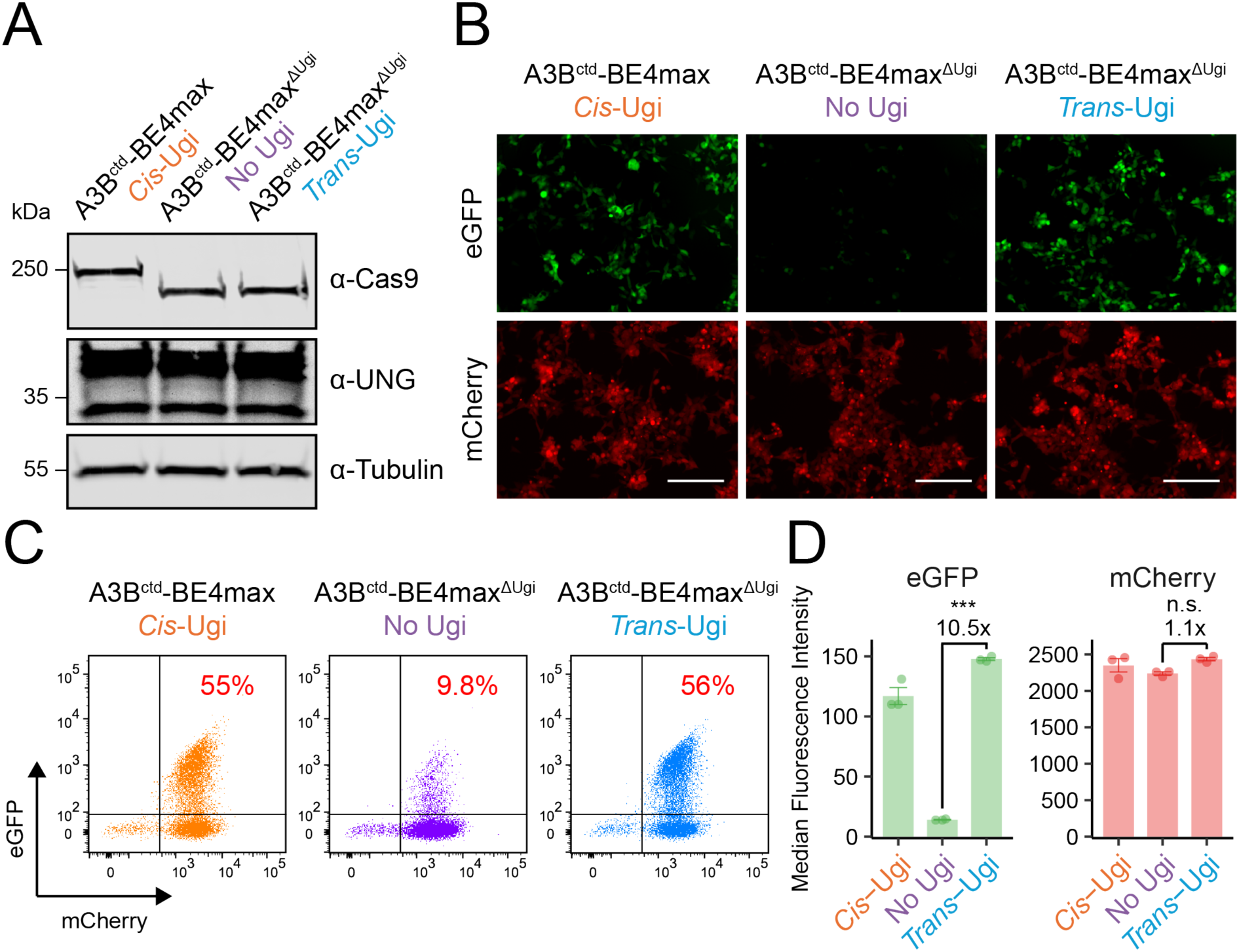
Chromosomal U-report system is sensitive to UNG activity. **A)** Immunoblot of Cas9 and UNG in the tested conditions 48 hours post-transfection of the 293T-chromosomal AMBER c15 cells. **B)** Fluorescence microscopy of cells from panel-A showing that eGFP levels correlate with UNG inhibition (*cis* or *trans*) and that mCherry expression remains stable at 48 hours post-transfection (scale=200μm). **C-D**) Flow analysis and quantification of the singlet population at the experimental end point. The percentages in red indicate the ratio double-positive to mCherry-positive cell populations at 48 hours post-transfection (*n*=3, mean±SEM).

### *UNG2* ablation is insufficient for full inhibition of UNG activity in the nucleus

The human genome encodes two variants of UNG, the mitochondrial UNG1 and nuclear UNG2 isoforms. The expression of each variant is driven by alternative transcription initiation sites and alternative splicing, leading to isoforms with distinct N-terminal sequences that dictate subcellular localization and function [13]. Given that the U-report biosensor, in both episomal and chromosomal formats, is designed to detect UNG activity in the nucleus, we hypothesized that our assay should only be sensitive to UNG2 activity. To test this, CRISPR was used to knockout (KO) *UNG1*, *UNG2*, and both variants simultaneously by targeting of isoform-specific or shared regions of the *UNG* gene on chromosome 12 (**Fig. 4A**). Independent clones were validated for each KO in 293T cells by immunoblotting and by uracil excision assays with whole cell extracts (**Fig. 4B**). Genetic inactivation was further confirmed by deep and Sanger sequencing the regions spanning each CRISPR cleavage site (**Supplementary Table S2**).

**Fig. 4.**
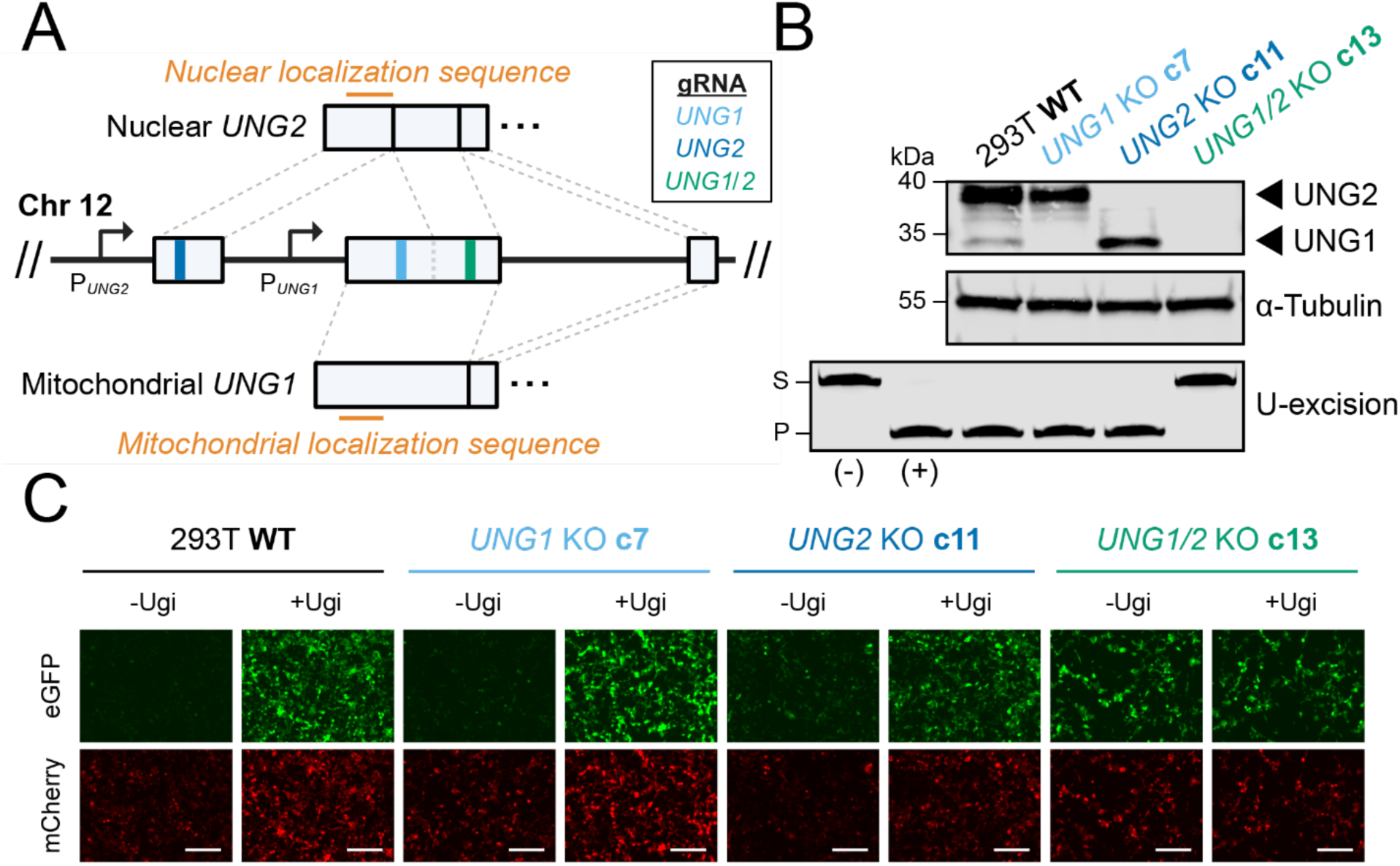
Inactivation of both mitochondrial *UNG1* and nuclear *UNG2* is required for full inhibition of UNG activity in the nucleus. **A)** Schematic of the *UNG* locus in the human genome and the regions targeted for disruption with CRISPR/Cas9. **B)** Validation of *UNG* knockouts in 293T cells by immunoblotting and *in vitro* uracil excision assay [S, uracil-containing 43-mer ssDNA substrate; P, cleaved 30-mer ssDNA product; (-) = HED buffer only (negative control); (+) = 5 Units of *E. coli* uracil-DNA glycosylase (positive control for uracil excision)]. **C)** Fluorescence microscopy at 48 hours post-transfection showing that *UNG1/2* double-KO is necessary for full inhibition of nuclear UNG activity (scale=200μm; corresponding flow cytometry analysis in **Supplementary Figure S5C**).

We next deployed the U-report biosensor in wildtype and KO cells by co-transfection and compared the fluorescence intensity between the absence and over-expression of Ugi. As predicted, the *UNG1* KO alone was insufficient to fully inhibit the system, as exhibited by low and high eGFP in the no and over-expressed Ugi conditions, respectively (**Fig. 4C**). Interestingly, although *UNG2* KO without Ugi led to a greater level of eGFP compared to wildtype without Ugi, the addition of excess Ugi in the *UNG2* KO raised the eGFP signal even higher (**Fig. 4C**). These results suggested that some capacity for UNG activity in the nucleus remained after *UNG2* disruption. Indeed, only the *UNG1/2* double KO caused full inhibition to be reached as indicated by equal eGFP activities with and without Ugi over-expression (**Fig. 4C**). These results were reproduced in a second set of 293T *UNG* KO clones (**Supplementary Fig. S5A-B**). Moreover, flow cytometry analysis of the eGFP-to-mCherry MFI ratio is consistent with the findings from fluorescence imaging of both sets of clones (**Supplementary Fig. S5C**).

To determine whether these results can be generalized beyond 293T cells, a systematic knockout approach was also used to disrupt each *UNG* isoform in U2OS cells (immunoblots in **Supplementary Fig. S6A**; genotypes in **Supplementary Table S2**). As above for 293T, neither the *UNG1* nor the *UNG2* knockouts were individually sufficient to ablate UNG activity in whole cell extracts (uracil excision data in **Supplementary Fig. S6B**). Only the *UNG1/2* double KO proved sufficient to completely ablate UNG activity in whole cell extracts (**Supplementary Fig. S6B**). U-report experiments were also performed as above but, despite multiple attempts, the transfection efficiencies were considerably lower overall for U2OS cells and were more variable between the individual clones. Limited delivery of the U-report biosensor components (*i.e*., AMBER, A3B^ctd^-BE4max^ΔUgi^, gRNA, and Ugi or empty plasmid control) constrains the degree of eGFP activation and, in turn, the sensitivity of the assay to UNG activity and its inhibition. These transfection issues resulted in less pronounced changes in the eGFP/mCherry MFI ratios between the no Ugi and excess Ugi conditions, with only UNG1 KO clones significantly affected by Ugi over-expression (fold-change in MFI ratio > 1; **Supplementary Fig. S6C**). Nonetheless, the *UNG2* KO clones also showed a fold-change in MFI ratio > 1 with excess Ugi consistent with a modest level of UNG1 activity in the nucleus, whereas the *UNG1/2* double KO exhibited a MFI ratio < 1 (**Supplementary Fig. S6C**). Together, these U2OS knockout experiments further indicated that both UNG1 and UNG2 contribute to the modulation of the U-report biosensor activity in the nucleus of living cells.

## Discussion

Here, we engineer a U-report biosensor capable of quantifying UNG activity in the nuclear compartment of living cells. This biosensor combines a modified cytosine base editor lacking Ugi proteins (A3B^ctd^-BE4max^ΔUgi^) and the AMBER fluorescent reporter system that is sensitive to C-to-U base changes [33,35]. The U-report system is responsive to a broad range of UNG activity. Systematic investigation of the endogenous UNG activity in human cells showed that both the mitochondrial UNG1 and nuclear UNG2 isoforms are active in the nucleus. One explanation for the nuclear activity of UNG1 is that its small size (∼34 kDa) may allow it to diffuse passively across the nuclear pore complex, if it is not being trafficked actively into mitochondria [40,41]. Alternatively, the action of UNG1 on genomic DNA could be cell cycle-dependent, as breakdown of the nuclear envelope during mitosis may expose nuclear DNA to UNG1 from the cytosol. Further studies are needed to fully elucidate the biological role of UNG1 activity in the nucleus its relationship to UNG2.

Our studies are consistent with a prior report showing that UNG1 can support class switch recombination in the mouse B cell lymphoma line CH12F3 [42]. CH12F3 cells express UNG2 and two variants of UNG1, an unprocessed high-molecular-weight (HMW) form and a processed low-molecular-weight (LMW) form. UNG1-HMW was observed in the nucleus of CH12F3 cells through cellular fractionation. *UNG* isoform-specific knockout clones of human cell lines (L428, HAP1, and U373) were generated and used for DNA uracil quantification by liquid chromatography-tandem mass spectrometry. The results of these studies also indicated that UNG1 contributes to nuclear uracil excision in the absence of the UNG2 isoform [42]. However, because cell lysis is an essential step toward genomic DNA purification and mass spectrometry analysis, it is possible that cytoplasmic UNG1 could have transiently contacted genomic DNA during sample processing. Our findings with the U-report system indicate that this is unlikely to be the case and that UNG1 exhibits a robust uracil excision activity in the nuclear compartment of living cells. During development of the U-report biosensor, we noticed that the mCherry fluorescence levels change in accordance with eGFP levels in transient transfection experiments. This is likely due to foreign DNA restriction, which has been described by our group and others [37–39]. The mechanism involves, first, plasmid DNA deamination by the cytosine base editor. The uracil intermediates then serve as substrates for UNG-mediated uracil excision. The resulting abasic sites are converted to DNA breaks during subsequent UBER steps, which in aggregate can destabilize the transfected plasmid. This issue was circumvented by integrating the AMBER construct into the chromosome for stable expression. The stable AMBER approach resulted in uniform expression of mCherry from the genome, and it improved signal-to-noise levels in the U-report biosensor (∼10x chromosomally versus ∼8x episomally).

A previously reported plasmid-based assay for measuring UNG activity in cells utilizes the conversion of a pre-engineered U:G mismatch to a C:G base pair in *BFP* to modulate blue fluorescence [32]. Although similarities with the U-report biosensor include a reliance on transfection and a fluorescence readout, there are three key differences. First is the need to pre-install a U:G mismatch in the plasmid construct, versus *de novo* creation of the uracil inside the nuclear compartment of the cell by APOBEC-catalyzed C-to-U deamination of an *eGFP* target site in the AMBER reporter. Second, the U-report biosensor encodes a constitutively expressed mCherry marker, which controls for transfection efficiency (episomal), strength of expression (chromosomal), and construct stability (episomal and chromosomal). The third and most important difference is that the U-report system can be utilized following integration into chromosomal DNA. This is particularly advantageous for examining UNG activity on a chromosome-like substrate under a wide range of genetic and/or pharmacologic conditions. Together, our studies demonstrate a powerful new approach for quantifying UNG activity in the nuclear compartment of living cells, and they also help to advance our collective understanding of UNG isoform functionality.

## Acknowledgements

We thank members of the Harris laboratory and Drs. Daniel Salamango and Arun Wiita for support and constructive feedback. BioRender was used to create the **Graphical Abstract** and **Fig. 1A**, **Fig. 1B**, **Fig. 1C**, and **Fig. 4A**.

## Supplementary data

Supplementary Data are available at NAR Online.

