## Supplementary Tables S1-S2 and Supplementary Figures S1-S6 for "Genetic studies with a uracil DNA glycosylase biosensor support a role for mitochondrial UNG1 in nuclear uracil repair"

**Supplementary Figures S1-S6**

### Supplementary Table S1. Oligonucleotide sequences.

| Oligo name | Purpose | Sequence 5' to 3' |
|---|---|---|
| hUNG2-insert-cloning-F | Forward PCR primer for amplifying hUNG2 followed by restriction cloning into pcDNA3.1 | CTTAAGCTTGCCACCATGATCGGCCAGAAG |
| hUNG2-insert-cloning-R | Reverse PCR primer for amplifying hUNG2 followed by restriction cloning into pcDNA3.1 | GCAGAATTCCCTACAGCTCCTTCCAG |
| remove-Ugi-SDM-F | Forward SDM primer for removing Ugi's from A3Bctd-BE4max | GAGGTGACTCTGGCGGCTCAAAA |
| remove-Ugi-SDM-R | Reverse SDM primer for removing Ugi's from A3Bctd-BE4max | CCGCCAGAGTCACCTCCCAGCTG |
| Ugi-insert-F | Forward PCR primer for amplifying Ugi insert for InFusion cloning | GGATCCGCCACCATGACTAATCTGAGCGACATC |
| Ugi-insert-R | Reverse PCR primer for amplifying Ugi insert for InFusion cloning | TCTGCAGAATTCTTACAGCATCTTGATCTTATTC |
| Ugi-backbone-F | Forward PCR primer for amplifying pcDNA3.1 backbone for InFusion cloning | CATGGTGGCGGATCCGAGCTCGGT |
| Ugi-backbone-R | Reverse PCR primer for amplifying pcDNA3.1 backbone for InFusion cloning | TAAGAATTCTGCAGATATCCAGC |
| Ugi-FLAG-SDM-F | Forward SDM primer to insert FLAG sequence into C-terminus of Ugi | GACTACAAGGACGACGATGACAAGTA-AGAATTCTGCAGATATCCAGCACAGTGG |
| Ugi-FLAG-SDM-R | Reverse SDM primer to insert FLAG sequence into C-terminus of Ugi | CTTGTCATCGTCGTCCTTGTAGTCCAG-CATCTTGATCTTATTCTCGCCGTTAG |
| hUNG2-PCR-F | Forward primer for generating amplicon from gDNA to confirm human *UNG2* KO | CACTGGGCCTTCCACCG |
| hUNG2-PCR-R | Reverse primer for generating amplicon from gDNA to confirm human *UNG2* KO | CTCTGATTGGACAGCGCACC |
| hUNG1-PCR-F | Forward primer for generating amplicon from gDNA to confirm human *UNG1* and *UNG1/2* KO | GGTGCGCTGTCCAATCAGAG |
| hUNG1-PCR-R | Reverse primer for generating amplicon from gDNA to confirm human *UNG1* and *UNG1/2* KO | TGGCCGGCTACACTAACAAG |
| remove-Ugi-sequencing | Sequencing primer to confirm Ugi removal from A3Bctd-BE4max | GCGATGCAATTTCCTCATTT |
| pcDNA3.1-sequencing | Sequencing primer to confirm inserts in pcDNA3.1 vectors | CGCAAATGGGCGGTAGGCGTG |
| pJET12F-sequencing | Sequencing primer to confirm inserts in pJET vectors | CGACTCACTATAGGGAGAGCGGC |

**Supplementary Table S2. *UNG* genotypes of knockout clones*.**

| Cells | Coding mRNA | Protein | Mutant allele 1 | Mutant allele 2 | Mutant allele 3 |
|---|---|---|---|---|---|
| 293T *UNG1* KO c7 | NM_003362.4 (*UNG1*) | NP_003353.1 (*UNG1*) | c.25_38del/p.(Trp9GlufsTer120) [2,739 reads] | c.30del/p.(Leu11TrpfsTer16) [2,601 reads] | |
| 293T *UNG1* KO c30 | NM_003362.4 (*UNG1*) | NP_003353.1 (*UNG1*) | c.25_38del/p.(Trp9GlufsTer120) [5,331 reads] | | |
| 293T *UNG2* KO c11 | NM_080911.3 (*UNG2*) | NP_550433.1 (*UNG2*) | c.3_5del/p.0 [1,664 reads] | c.[3_5del;32del]/p.0 [344 reads] | c.2_132+218delinsGCTG/p.0 [Sanger] |
| 293T *UNG2* KO c12 | NM_080911.3 (*UNG2*) | NP_550433.1 (*UNG2*) | c.2_7del/p.0 [2,496 reads] | c.-9_1del/p.0 [2,392 reads] | c.1_2ins[468bps]/p.0 [Sanger] |
| 293T *UNG1/2* KO c13 | NM_080911.3 (*UNG2*) | NP_550433.1 (*UNG2*) | c.262_272delinsA/p.(Arg88ThrfsTer24) [2,696 reads] | c.264_274del/p.(Asn89GlyfsTer50) [2,545 reads] | |
| 293T *UNG1/2* KO c52 | NM_080911.3 (*UNG2*) | NP_550433.1 (*UNG2*) | c.237_275delinsGGGGG/p.(Ala80GlyfsTer24) [5,291 reads] | c.273dup/ p.(Val92ArgfsTer51) [Sanger] | c.237_274del/ p.(Ala80GlyfsTer50) [Sanger] |
| U2OS *UNG1* KO c6 | NM_003362.4 (*UNG1*) | NP_003353.1 (*UNG1*) | c.25_38del/p.(Trp9GlufsTer120) [3,529 reads] | | |
| U2OS *UNG1* KO c7 | NM_003362.4 (*UNG1*) | NP_003353.1 (*UNG1*) | c.27_30delinsTTCCACAAATGGGCGTCTTCTGCC/p.(Trp9CysfsTer25) [4,670 reads] | | |
| U2OS *UNG2* KO c2 | NM_080911.3 (*UNG2*) | NP_550433.1 (*UNG2*) | c.1_88del/p.0 [2,969 reads] | c.-1_12del/p.0 [2169 reads] | |
| U2OS *UNG2* KO c7 | NM_080911.3 (*UNG2*) | NP_550433.1 (*UNG2*) | c.-1_12del/p.0 [4,348 reads] | | |
| U2OS *UNG1/2* KO c3 | NM_080911.3 (*UNG2*) | NP_550433.1 (*UNG2*) | c.272_273insT/p.(Val92ArgfsTer51) [4,713 reads] | | |
| U2OS *UNG1/2* KO c8 | NM_080911.3 (*UNG2*) | NP_550433.1 (*UNG2*) | c.267_295del/p.(Asn89LysfsTer44) [2,098 reads] | c.241_274del/p.(Ala81TrpfsTer23) [2,039 reads] | |

* 293T and U2OS have abnormal karyotypes with 3 and 2 copies of the *UNG* locus, respectively. Based on our sequence analyses, clones with fewer alleles are likely due to identical CRISPR resolutions, loss-of-heterozygosity during CRISPR-mediated genome engineering, and/or to deletions that extend beyond the interval analyzed by PCR and sequencing (*i.e.*, large null alleles).

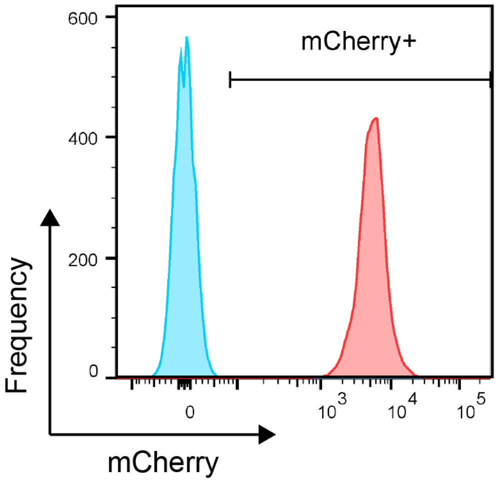


**Supplementary Figure S1. Flow cytometry analysis of 293T-cAMBER c15.**

Flow analysis of parental 293T cells (blue) and 293T-cAMBER c15 cells (red), which has the AMBER construct integrated stably into chromosomal DNA. The precise insertion site has yet to be identified but is likely single copy based on uniformly high mCherry expression levels.

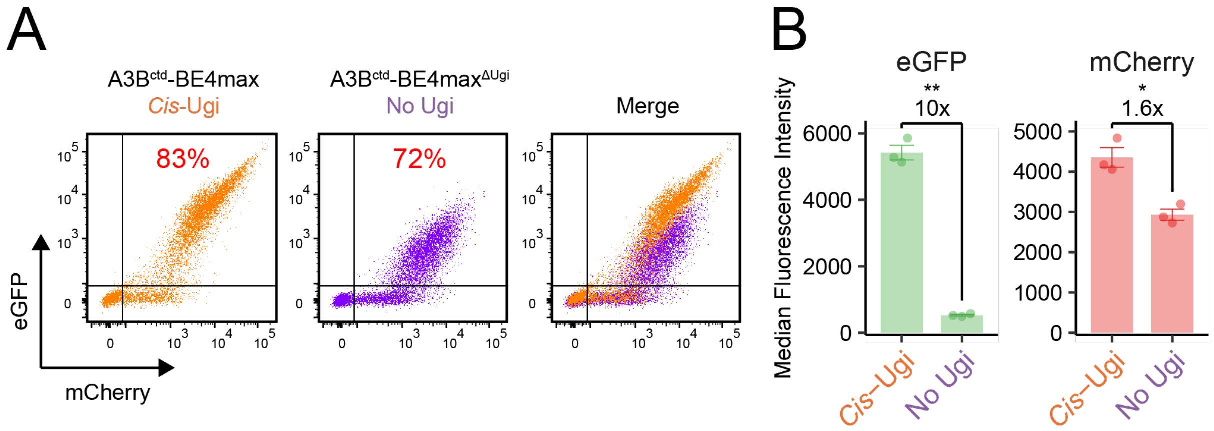


**Supplementary Figure S2. Flow cytometric confirmation of changes in fluorescence upon Ugi removal.**

**A-B**) Flow analysis and quantification of the singlet population harvested 48 hours post-transfection showing reduced eGFP and mCherry levels when Ugi is removed. The percentages in red indicate the ratio double-positive to mCherry-positive cell populations (*n*=3, mean±SEM).

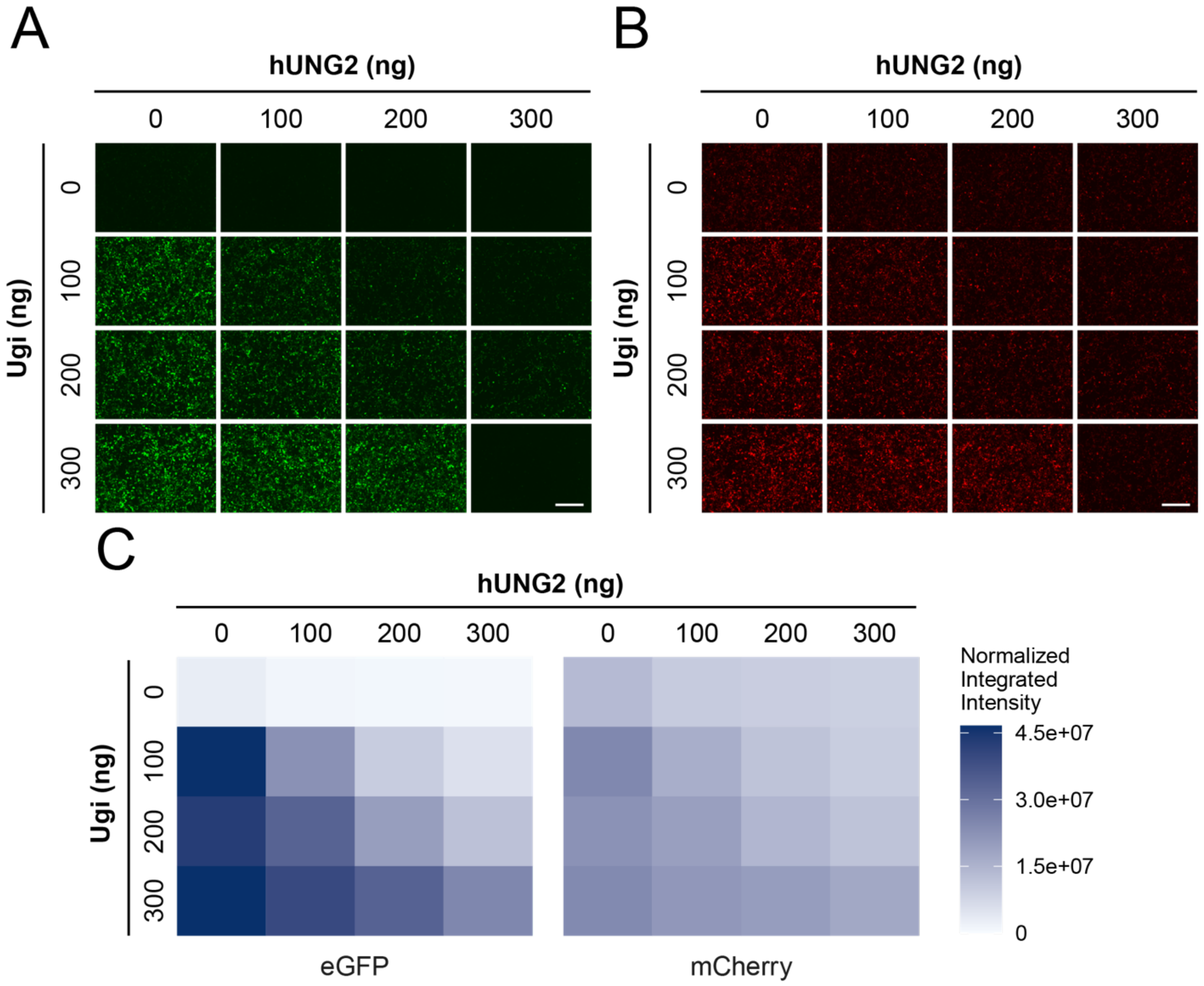


**Supplementary Figure S3. U-report biosensor is responsive to varying levels of UNG activity.**

**A-B**) Fluorescence microscopy of eGFP (left) and mCherry (right) signals 48 hours post-transfection of 293T cells with Ugi and human UNG2 (hUNG2) in the episomal system (scale=400µm).

**C**) Quantification of the fluorescence images at 48 hours post-transfection.

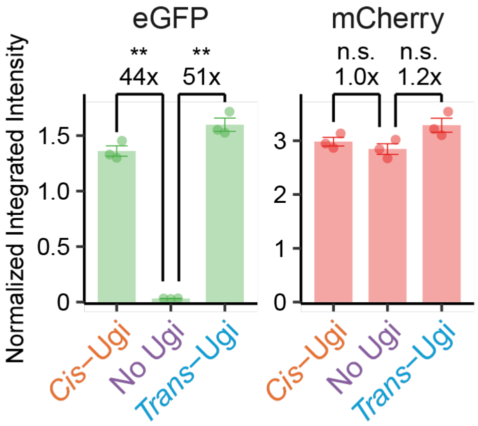


**Supplementary Figure S4. Quantification of fluorescence imaging of the chromosomal U-report system.**

Comparison of the normalized integrated intensities of eGFP and mCherry across the three conditions in the 293T-chromosomal AMBER c15 cells at 48 hours post-transfection (*n*=3, mean±SEM).

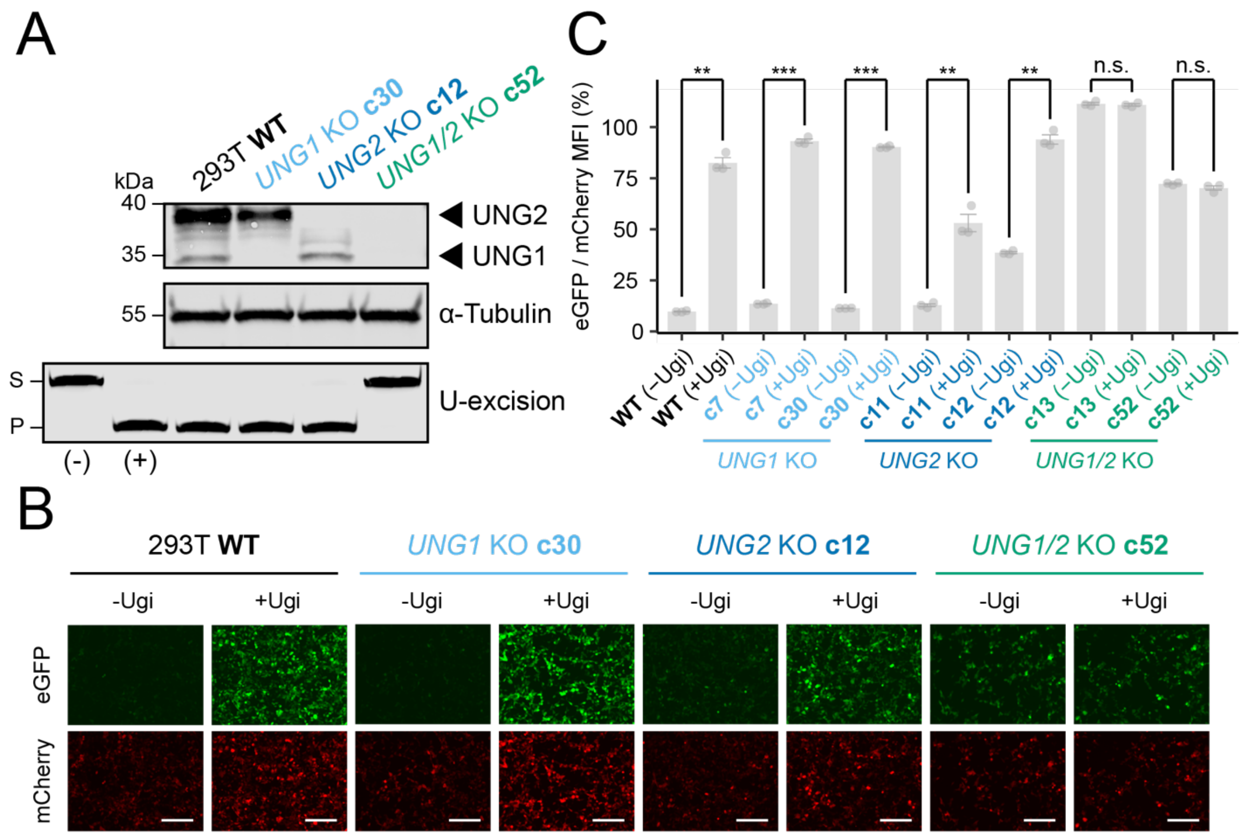


**Supplementary Figure S5. U-report biosensor in 293T *UNG* KO clones.**

**A**) Immunoblot and *in vitro* uracil excision assay confirming *UNG* inactivation in 293T cells [S, uracil-containing 43-mer ssDNA substrate; P, cleaved 30-mer ssDNA product; (-) = HED buffer only (negative control); (+) = 5 Units of *E. coli* uracil-DNA glycosylase (positive control for uracil excision)].

**B**) Fluorescence microscopy of additional *UNG* KO clones in 293T cells at 48 hours post-transfection (scale=200μm).

**C**) Ratio of eGFP to mCherry median fluorescence intensities at 48 hours post-transfection shows full UNG inhibition only in *UNG1/2* double-KO clones (*n*=3, mean±SEM). Quantification and statistical comparisons are only relevant, as indicated, for individual clones (minus/plus Ugi) due to the variability of transfection efficiencies between independent clones.

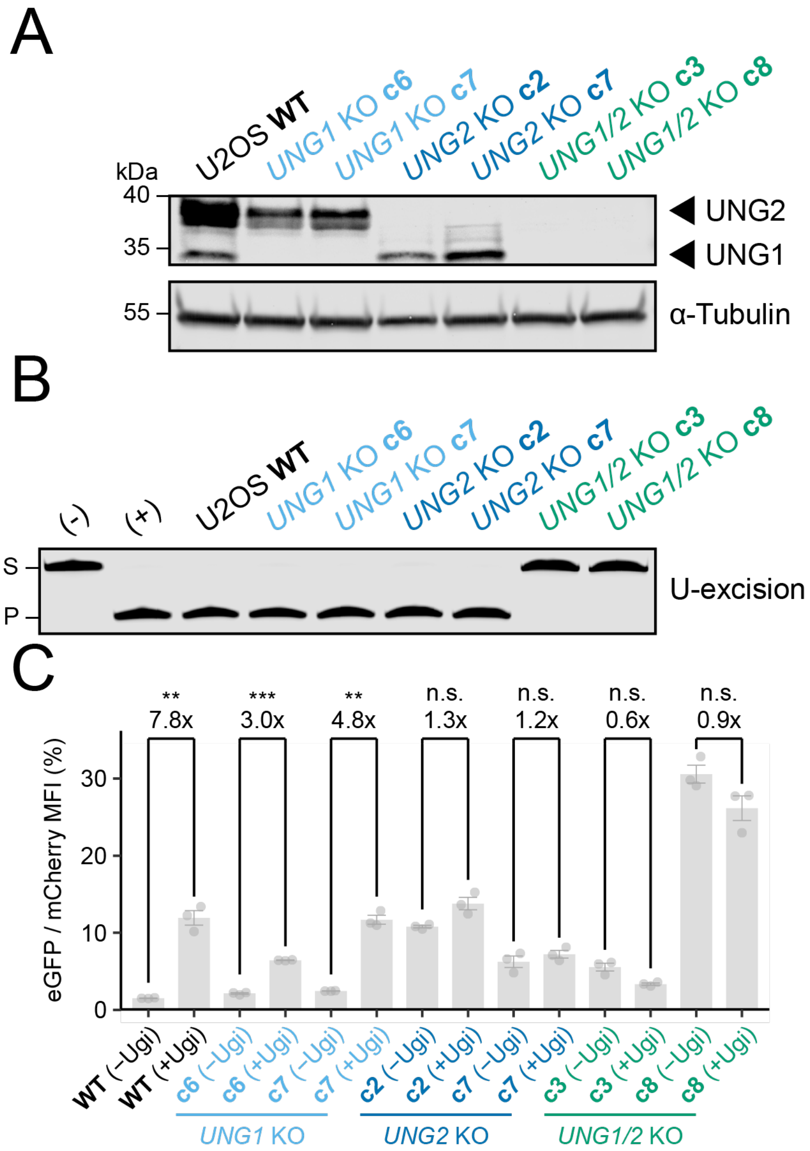


**Supplementary Figure S6. U-report biosensor in U2OS *UNG* KO clones.**

**A**) Immunoblot validation of *UNG* isoform disruption in U2OS cells.

**B**) Uracil excision activity in whole cell extracts from the indicated clones [S, uracil-containing 43-mer ssDNA substrate; P, cleaved 30-mer ssDNA product; (-) = HED buffer only (negative control); (+) = 5 Units of *E. coli* uracil-DNA glycosylase (positive control for uracil excision)].

**C**) Ratio of eGFP to mCherry median fluorescence intensities at 48 hours post-transfection using the U-report system. Fold changes in the eGFP-to-mCherry MFI between the presence and absence of Ugi are indicated (*n*=3, mean±SEM). Quantification and statistical comparisons are only relevant, as indicated, for individual clones (minus/plus Ugi) due to the variability of transfection efficiencies between independent clones.
